# Using cell line and patient samples to improve predictions of patient drug response

**DOI:** 10.1101/026534

**Authors:** Cheng Zhao, Ying Li, Zhaleh Safikhani, Benjamin Haibe-Kains, Anna Goldenberg

**Affiliations:** SickKids Research Institute, 686 Bay Street, Toronto, ON M5G 0A4, Canada; Department of Computer Science, University of Toronto, 40 St. George Street, Toronto, ON M5S 2E4; Princess Margaret Cancer Centre, University Health Network, 101 College Street, Toronto, ON M5G 1L7; Department of Medical Biophysics, University of Toronto, 101 College Street, Toronto, ON M5G 1L7

**Author notes:** co-last and corresponding authors.

**Keywords:** Drug response, cell line, patient therapy response, data integration, bortezomib, erlotinib, docetaxel, epirubicin

## Abstract

**Background:** Recent advances in high-throughput technologies have facilitated the profiling of large panels of cancer cell lines with responses measured for thousands of drugs. The computational challenge is now to realize the potential of these data in predicting patients’ responses to these drugs in the clinic.

**Methods:** We address this issue by examining the spectrum of prediction models of patient response: models predicting directly from cell lines, those predicting directly from patients, and those trained on cell lines and patients at the same time. We tested 21 classification models on four drugs, that are bortezomib, erlotinib, docetaxel and epirubicin, for which clinical trial data were available.

**Results:** Our integrative models consistently outperform cell line-based predictors, indicating that there are limitations to the predictive potential of *in vitro* data alone. Furthermore, these integrative models achieve better predictive accuracy and require substantially fewer patients than would be the case if only patient data were available.

**Conclusions:** The integration of *in vitro* and *ex vivo* genomic data results in more accurate predictors using only a fraction of the patient information, which can help optimize the development of personalized predictors of therapy response. Altogether our results support the relevance of preclinical data for therapy prediction in clinical trials, enabling more efficient and cost-effective trial design.

## Background

Developing molecular predictors of therapy response^1^ is the key to implementing precision medicine in the clinic. Such predictors would allow clinicians to select the best available therapeutic option for each individual patient. The classical approach to building drug response-predictive tests consists of correlating molecular profiles of patient tumors with drug response outcome data collected during clinical trials. In this context, previous studies used gene expression profiles of tumors at diagnosis to predict patients’ therapy response. Chang et al. investigated the predictive value of tumor gene expression profiles in neoadjuvant setting for advanced breast cancer patients treated with docetaxel ^1^. Mulligan et al. developed an expression-based predictor of response to bortezomib for patients with relapsed myeloma enrolled in phase II/III clinical trials ^2^. Kim et al. assessed the accuracy of four pre-specified genetic and transcriptomic biomarkers in the Biomarker-integrated Approaches of Targeted Therapy for Lung Cancer Elimination (BATTLE) trial in non – small cell lung cancer ^3^. In our previous work, we developed expression-based TOP2A amplification, tumor invasion and immune response signatures, yielding a high negative predictive value for response to epirubicin in breast cancer that does not express estrogen receptor ^4^. Though promising, the number of biomarkers identified through these and other studies used in clinical settings remains small ^5^. This is mainly due to the fact that most genomic predictors of drug response lack robustness and have not been validated in subsequent studies ^6^. The MicroArray Quality Control (MAQC) consortium showed that developing multivariate predictors of drug response in a complex disease like cancer is challenging ^7^. One of the limiting factors in many studies was the small size of the clinical cohort rather than the choice of the predictive approach ^1,3^. Unfortunately, it is not always feasible to collect large cohorts of patients that could improve the performance of the models ^8^. That said, these patient-based predictive models are still among the best predictors of drug response that we currently have and are the state-of-the-art in the clinic ^9^.

Recently, multiple studies have attempted to leverage molecular and pharmacological profiles of large panels of cancer cell lines in order to create genomic predictors of therapy response ^10–12^. These data are expected to boost the sample size and reduce the cost of building predictive models, while allowing pharmacological response screening for many drugs at a time ^10–12^. Such pharmacogenomic datasets were recently used to develop multivariate drug response-predictive models in non-small cell lung cancer (NSCLC), breast cancer and myeloma ^13,14^. However, using only cell lines to build predictive models of patient drug response presents its own difficulties. For example, it is well established that *in vitro* models, such as cancer cell lines, exhibit substantially different molecular features than patient tumors ^15,16^. This inconsistency often results in a poor validation accuracy of genomic predictors on independent cohorts of patients ^17^. Moreover, we recently showed that response labels for the same cell lines to the same drugs are not always consistent across different studies, potentially further hampering the reproducibility of cell line based predictors ^18–20^.

In the present work we aim to take maximal advantage of the existing *in vitro* and *ex vivo* data to improve prediction accuracy of drug response in patients. We show that predictive models that combine cell line and patient data during model training significantly improve upon existing work. By incorporating multiple data sources into the training set, we increase its sample size, but more importantly, we take into account similarity of cell lines and patient data during model development, allowing us to mitigate the inherent cell line-patient differences compared to cell line-based predictive approaches. To the best of our knowledge, this study describes the first computational pipeline efficiently combining *in vitro* and *ex vivo* samples to develop robust molecular predictors of drug response. Our novel integrative approach significantly improves drug response prediction in patients while using fewer patient samples for training than models based on patient data alone. Our results support the use of preclinical data to build more accurate predictive models of response enabling improved implementation of adaptive clinical trials ^21^.

## Results

To test the predictive power of *in vitro* and *ex vivo* data integration, we considered three types of models: models developed using cell lines only (C2P models), those based on patient tumors only (P2P models) and models combining cell lines and patient data during training (CP2P models). To make patient drug response predictions we considered seven different machine learning approaches: Support Vector Machines with linear *SVM lin* and radial basis function *SVM rbf* kernels; *Ridge*, *Lasso* and *Elastic Net* regressions; Random Forest *RF* and Similarity Network Fusion *SNF*. Recognizing the importance of feature selection for various predictive tools, we compared three feature selection techniques: all genes, 1000 landmark genes as defined in the LINCS project ^22^ and 1000 genes selected using the Minimum Redundancy, Maximum Relevance technique ^23^. More details are available in the Methods and Supplementary Methods.

Given the variance among different measures of cancer cell line sensitivity to drugs ^24,25^, we used three different binarized summary statistics of the drug dose-response curve, namely the drug concentration required to inhibit 50% of the maximal growth inhibition (IC_50_), the area under the dose-response curve (AUC) and the slope of the curve (Slope).

In total, we analyzed 504 possible model-scenario-outcome combinations across four drugs: bortezomib ^2^, erlotinib ^3^, docetaxel ^1^ and epirubicin ^4^, where both patients’ tumor gene expression profiles and their outcome data were available.

### Bortezomib in myeloma

Patients’ response to bortezomib were collected from the APEX phase 3, SUMMIT and CREST phase 2 trials with measured response to bortezomib in relapsed multiple myeloma patients ^2^. The microarray gene expression profiling was done on 169 samples collected from bone marrow aspirates, which were enriched for tumor cells (see Supplementary Tables S1 and S2 for more details). Cancer cell line drug response and microarray data were derived from the Cancer Genome Project (CGP) ^11^ where 313 cancer cell lines across 26 tissues were treated with bortezomib. We applied Surrogate Variable Analysis (SVA) ^26^ to homogenize cell line mRNA expression data and patient expression data (Figures 2A,B).

**Figure 1.**
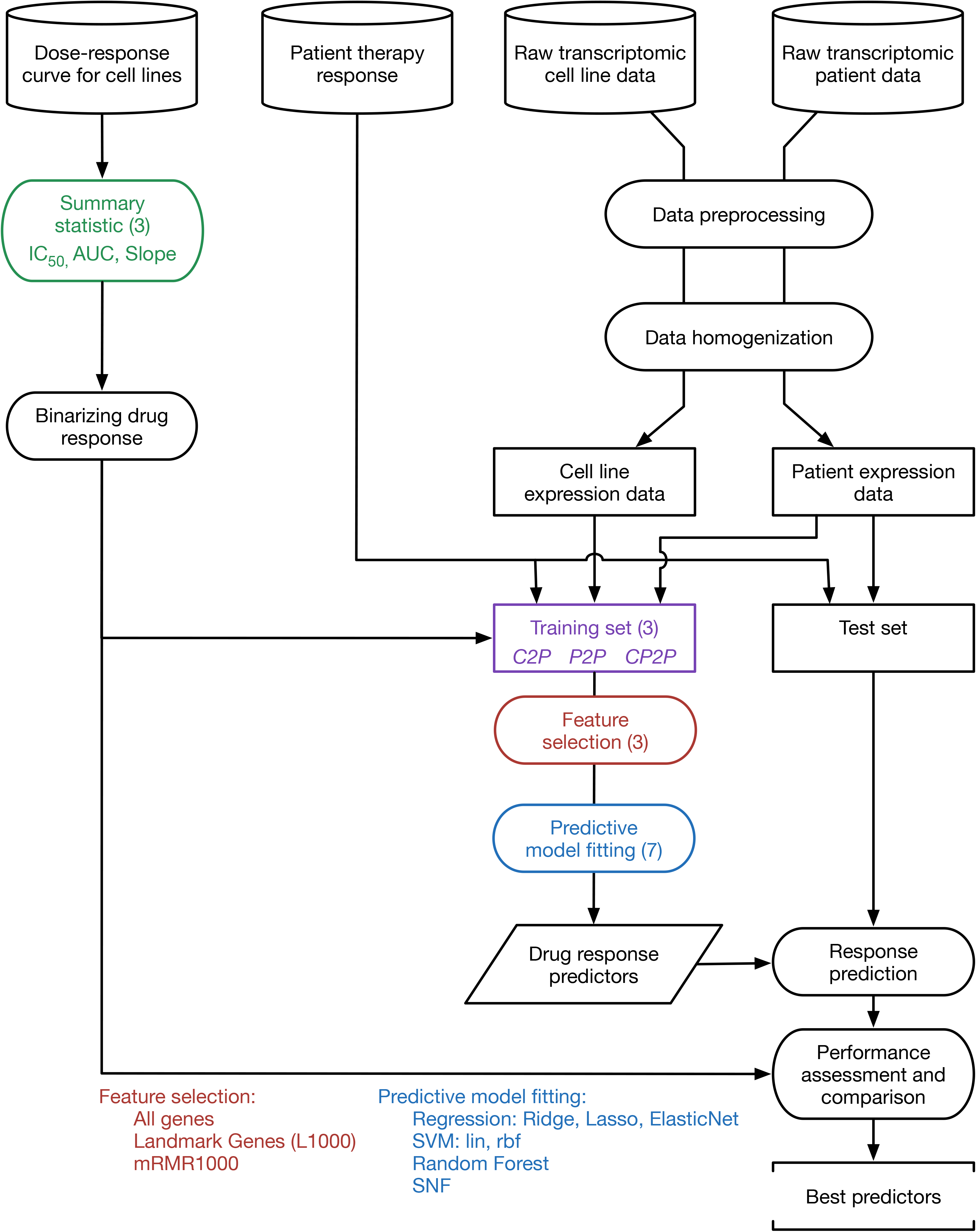
Schematic of our approach for prediction modeling of drug response.

**Figure 2.**
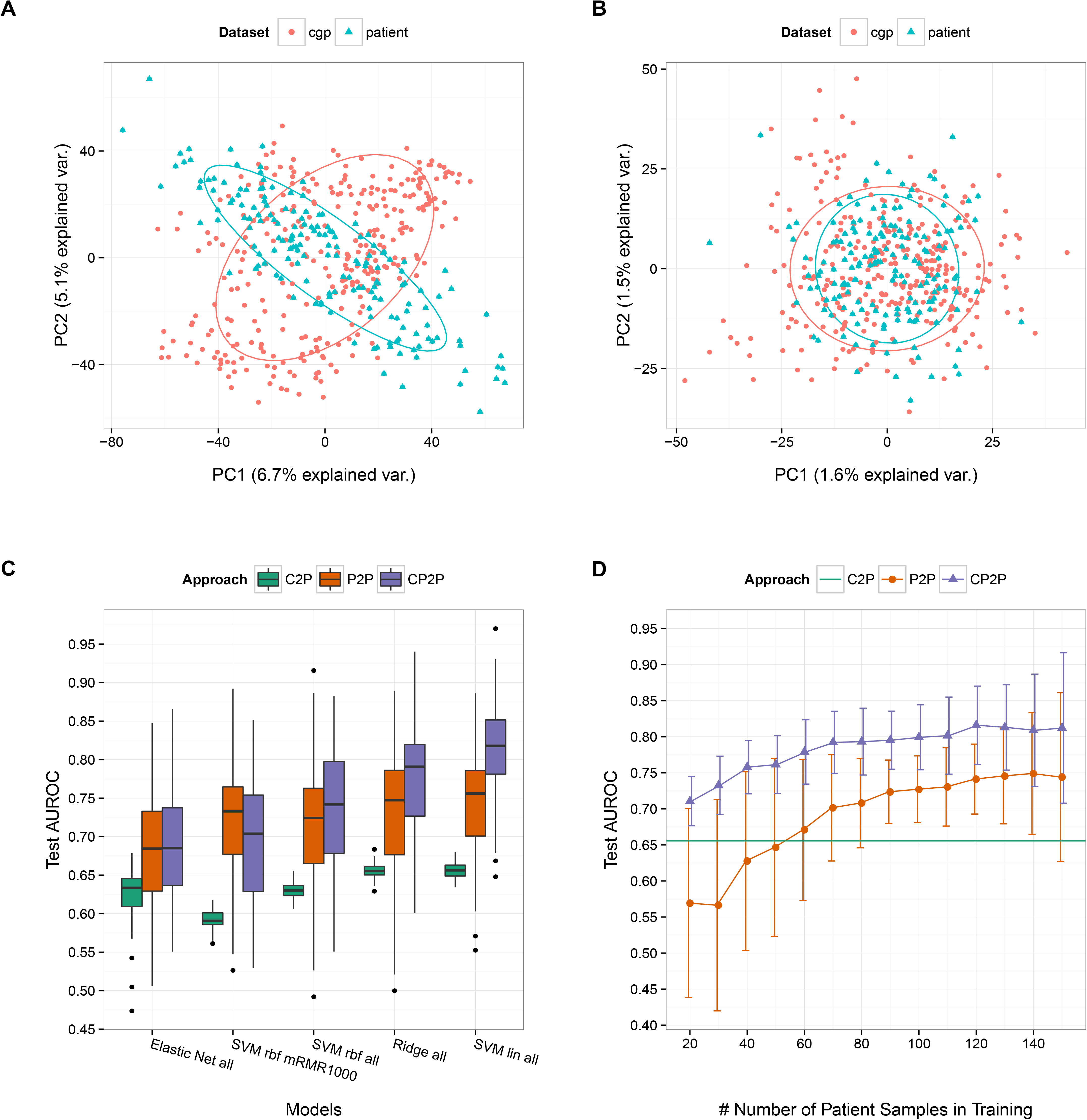
Results for bortezomib. (A) Plot of the first and second principal components for the patient and CGP datasets with no batch effect corrections. (B) Plot of the first and second principal components for the patient and CGP datasets when SVA with 2 surrogate variables is applied to homogenize the two datasets. The ellipses represent one standard deviation away from the mean by fitting the Gaussian to each data type. (C) The performance of the top 3 models for each approach are plotted using the Slope summary statistics, resulting in 5 distinct methods. For C2P, 300 cell line samples are used for training. For P2P, 130 patient samples were used for training. For CP2P, 311 cell line and 130 patient samples were used for training. (D) Comparison of the best P2P, C2P-Slope, and CP2P-Slope models with varying number of patient samples in training data.

#### Comparing models

We trained 147 predictive models (see Methods) using 5-fold cross-validation and assessed their performance (AUROC) on a held out set, repeating this procedure a 100 times. The top models combining cell line and patient tumor data (CP2P) yielded significantly higher predictive performance than the top C2P and P2P models (p = 2.2e-16 and 4.5e-07 using a paired one-sided Mann-Whitney test, Figure 2C). The best-performing model was an SVM with a linear kernel using gene expression of all genes (*SVM lin all*) with the binarized Slope of the drug-dose response curves as the pharmacological outcome. *SVM lin all* model performed significantly better than the next-best performer (*Ridge all*) when combining patients and cell lines during training (p = 2.91e-10 using a paired one-sided Mann-Whitney test). In P2P and C2P settings, there was no statistically significant difference between *SVM lin all* and *Ridge all* models, yet both of these methods were significantly better than the others (Figure 2C). Among the outcome measures, the best predictive method used binarized Slope as drug response outcome in cell lines (median AUROC = 83%, 77% and 76% for Slope, AUC and IC_50_, respectively; Supplementary Table S3). However, the Slope summary statistic was not the best in all settings. For example, in the C2P-AUC setting, when 300 cell line samples were used to train the models, *Ridge mRMR1000* achieved a test AUROC of 68% (Supplementary Figure S3), performing significantly better than the best model for C2P-Slope (p = 2.2e-16 using a paired one-sided Mann-Whitney test). Concurring with our previous study ^18^, binarized IC**_50_**seemed to be the worst summary statistic to build predictors of bortezomib response and Slope was the best (Supplementary Table S3), with the most drastic difference in AUROC of 21% between CP2P-Slope and C2P-IC50.

#### Power analysis

We observed that the best CP2P models were consistently outperforming the best C2P models. We then assessed the minimum number of patients that in addition to cell lines (at training) could achieve the same or better accuracy than using just patient data. Figure 2D shows the comparison of P2P, C2P, and CP2P as we vary the number of patients. When the number of patients is small, the best cell line-based classifiers significantly outperform the best patient-based classifiers (one-sided Mann-Whitney test p-value < 3.5e-4). The best patient-based classifier performs as well as or better than the best C2P classifier when at least 40 patients are used for training. The best CP2P model performed better than the best C2P and P2P models for the full range of patient numbers, needing as few as 60 patients to outperform the best P2P classifier with 150 patients (p = 0.012 using a one-sided Mann-Whitney test). Our results suggest that using our framework it is possible to match the performance of patient-based predictive models while recruiting far fewer patients.

### Erlotinib in non-small cell lung cancer

We retrieved tumor gene expression profile and therapy response from the Biomarker-Integrated Approaches of Targeted Therapy for Lung Cancer Elimination (BATTLE) trial ^3^. A subset of 25 patients with recurrent or metastatic non-small cell lung cancer (NSCLC) were treated with erlotinib. Therapy response was defined as progression-free survival time of 2 or more months. For *in vitro* data, we used 287 cell lines from CGP (41 lung cancer) and 44 NSCLC cell lines from the BATTLE study with IC_50_ drug response values only (Supplementary Tables S4-5). The three datasets were homogenized using ComBat ^27^ (Figure 3A). The homogenization was improved when all CGP cell lines across tissues were used (Figure 3B).

**Figure 3.**
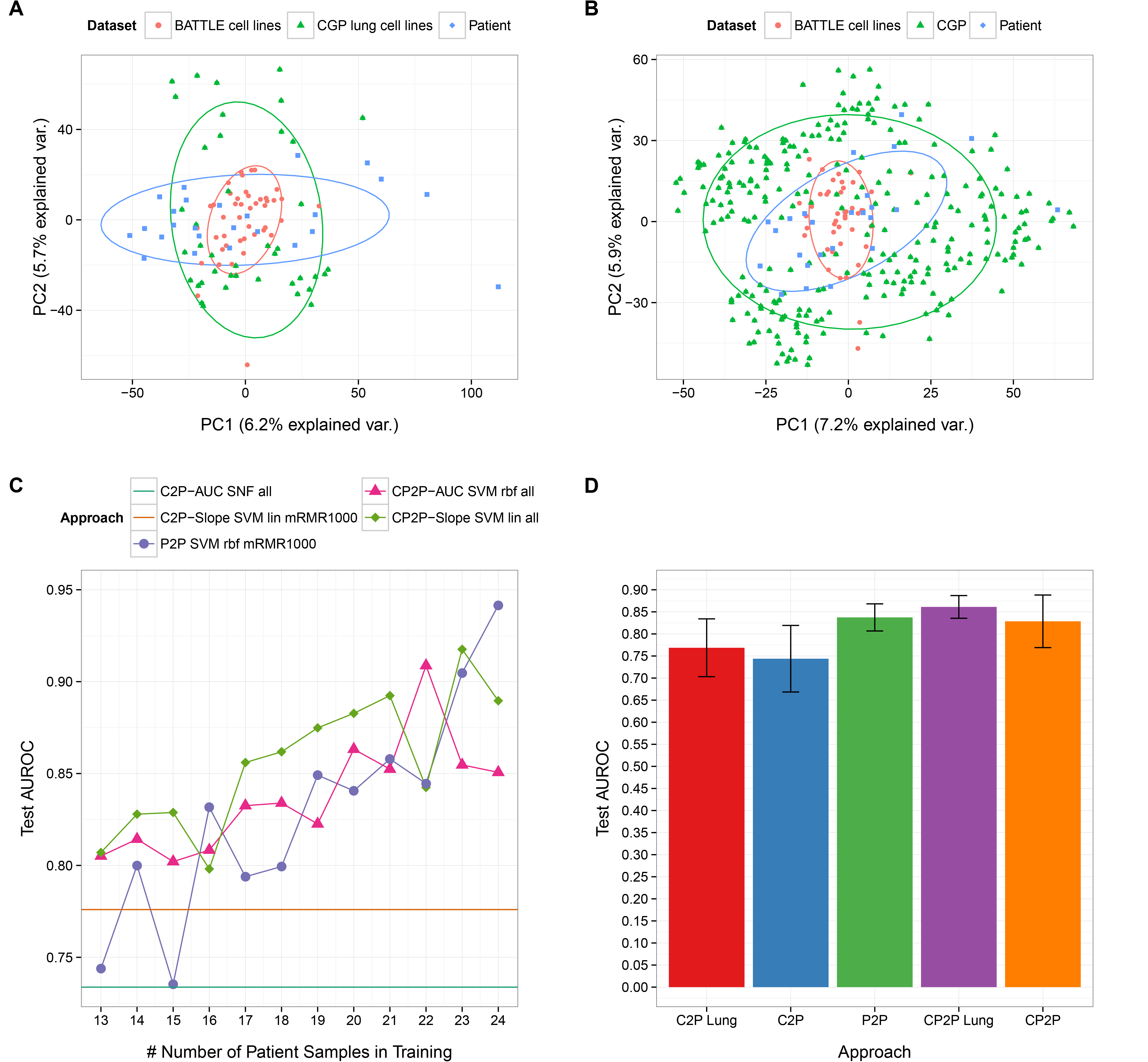
Results for erlotinib. (A) Plot of the first and second principal components for the patient dataset and lung cell lines from the CGP and BATTLE datasets when ComBat is applied to homogenize the three datasets. (B) Plot of the first and second principal components for the patient, all of the CGP cell lines tested with erlotinib, and BATTLE datasets when ComBat is applied to homogenize the three datasets. The ellipses represent one standard deviation away from the mean by fitting the Gaussian to each data type. (C) Comparison of the performance for the best model from each approach using different summary statistics with varying number of patient samples in the training set. IC_50_ and AUC summary statistics produced identical labels and therefore only one of them is plotted. (D) Comparison of the performance for the best model using AUC summary statistics from each approach. Models trained with only lung cancer cell lines are also compared. For C2P Lung, 80 lung cell line samples were used for training. For C2P, 300 cell line samples used for training. For P2P, 13 to 24 patient samples were used for training. For CP2P Lung, 85 lung cancer cell lines and 13 to 24 patient samples were used for training. For CP2P, 331 cell line and 13 to 24 patient samples were used. The mean and standard deviation over these ranges are shown.

Given that the erlotinib trial had few patients compared to the bortezomib trial, the significance analysis of the performance was based on varying the patient training set from 13 to 24 patients. The best CP2P model again outperformed the best C2P and P2P models with as few as 14 patients (Figure 3C). Interestingly, *SVM*s were still among the best performing models, though the kernel and the feature selection had larger impact on performance than in the case of bortezomib. Starting with 16 patients, the best C2P models were outperformed by both P2P and CP2P best models. Both P2P and CP2P models improved their AUROC with more patients in the training set, however performance of these P2P and CP2P models did not statistically differ. This could be due to the small number of patients and challenging homogenization of cell lines. It is interesting to note that using lung cancer cell lines to train the models produced more robust results (lower variance) but the same median performance compared to using all cell lines across tissues (Figure 3D).

### Docetaxel in breast cancer

The clinical dataset for docetaxel consisted of 24 patient samples with microarray gene expression obtained from breast cancer tumor biopsies before treatment ^1^. Response to docetaxel neoadjuvant treatment was based on whether 25% of the tumor remained after four cycles of docetaxel. We used 618 CGP cell lines, of which 40 were of breast tissue type and all were treated with docetaxel. The patient and cell line datasets were homogenized with ComBat (Figures 2A,B).

The docetaxel trial, like the erlotinib trial, contained fewer patients than the bortezomib trial. We therefore used varying number of patients to assess the significance in performance among the various methods (from 14 to 23 patients). The best CP2P model significantly outperformed the best C2P and P2P approaches when using either AUC or IC_50_ response summary statistics (p < 9.8e-04 for each using one-sided paired Mann-Whitney tests) (Figure 4C). However, CP2P-Slope performance did not significantly differ from the P2P performance (p = 0.12 using a one-sided Mann-Whitney test). Interestingly, C2P performed better than P2P models for the AUC and IC_50_ statistics throughout the range of patient samples used for training. This result is not consistent with the case of erlotinib, where C2P classifiers were not as accurate and point to the apparent variance among patients responses to drugs. These results also indicate that there may not be one preferable summary statistic for the drug dose-response curves across different drugs.

**Figure 4.**
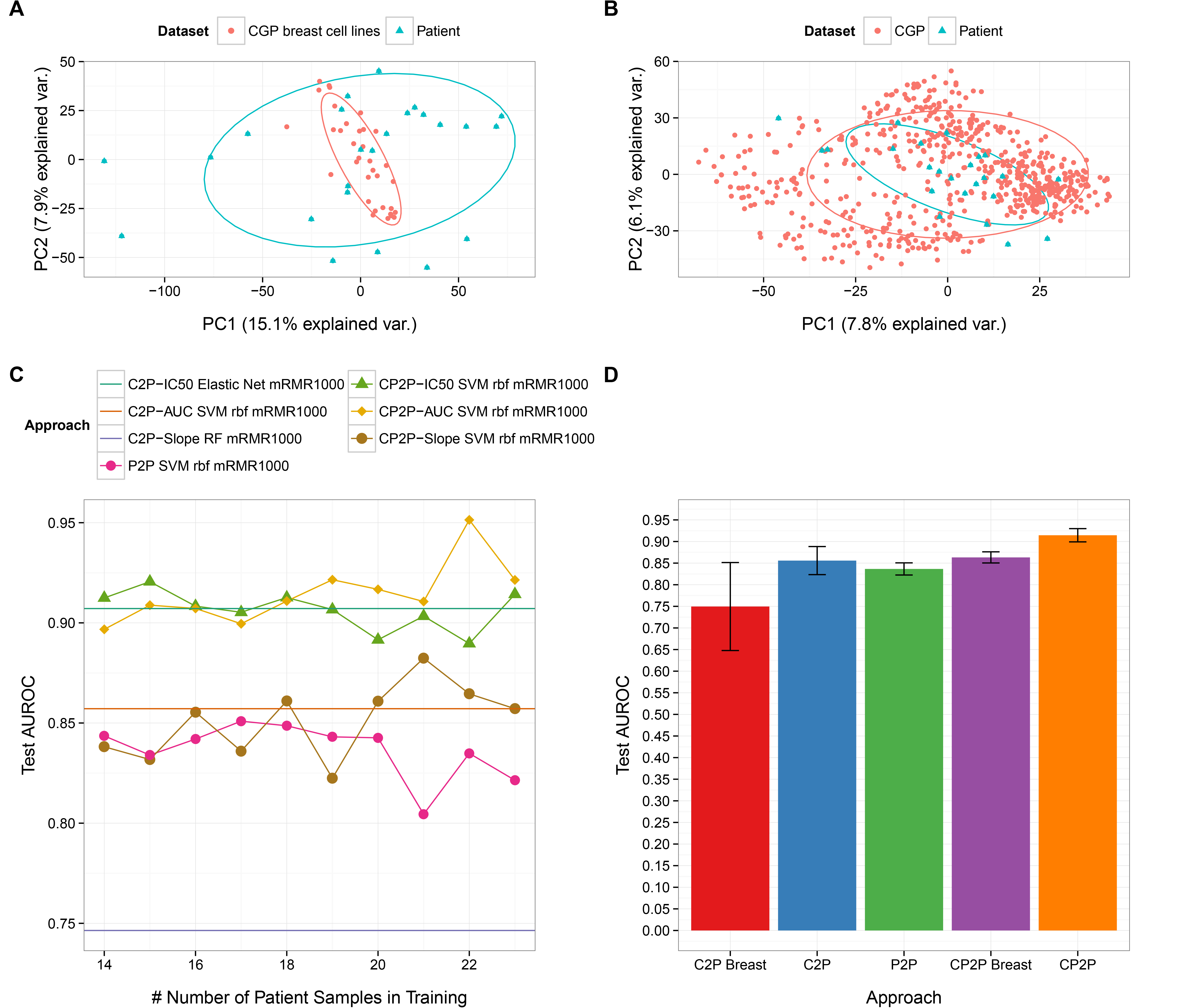
Results for docetaxel. (A) Plot of the first and second principal components for the patient dataset and breast cancer cell lines from CGP dataset homogenized using ComBat. (B) Plot of the first and second principal components for the patient and all cell lines from the CGP datasets homogenized using ComBat. The ellipses represent one standard deviation away from the mean by fitting the Gaussian to each data type. (C) Comparison of the performance for the best model from each approach with varying number of patient samples in the training set. (D) Comparison of the performance of the best methods using AUC summary statistics from each approach. Models trained with only breast cell lines are also compared. For C2P Breast, 30 breast cancer cell lines were used for training. For C2P, 580 cell lines were used for training. For P2P, 14 to 23 patient samples were used for training. For CP2P Breast, 34 breast cancer cell lines and 14 to 23 patient samples were used for training. For CP2P, 618 cell lines and 13 to 24 patient samples were used. The mean and standard deviation over the range of patient samples used for training are plotted.

We analyzed the performance with respect to the origin of cell lines^2^. For most scenarios, using only breast cell line samples did worse than using all cell lines. For CP2P-AUC, using all cell lines did significantly better than using breast cancer cell lines alone (p = 5.8e-16 using a one-sided Mann-Whitney test, Figure 4D). These results suggest that when the number of cell lines of the same tissue type as the patient’s cancer is small, using all cell lines may be preferable.

### Epirubicin in estrogen receptor-negative breast cancer

The clinical dataset for this set of experiments came from the neoadjuvant Trial of Principle (TOP) study, in which 118 patients with estrogen receptor-negative tumors were treated with epirubicin monotherapy ^4^. Patients were evaluated for pathologic complete response (pCR). The microarray gene expression profiling was done on samples collected from pre-treatment biopsy. No cell line samples in CGP were evaluated for response to epirubicin, and therefore we used 38 breast cancer cell lines from Heiser et al. ^28^. Note that in this study the number of cell lines is much smaller than the number of patients. We used SVA to homogenize patients with the cell line datasets (Figure 5A,B).

**Figure 5.**
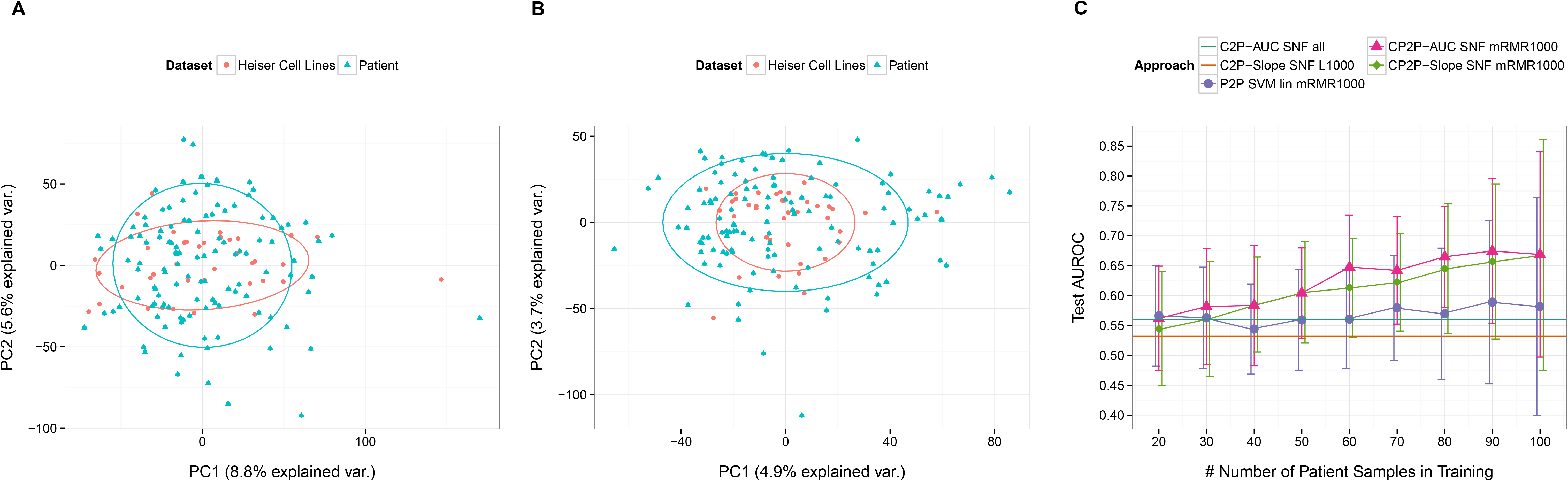
Results for epirubicin. (A) Plot of the first and second principal components for the original patient and Heiser data with no batch effect corrections. (B) PCA plot of the first and second principal components for the patient and Heiser datasets when SVA with 3 surrogate variables is applied to homogenize the two datasets. The ellipses represent one standard deviation away from the mean by fitting the Gaussian to each data type. (C) Comparison of the best models for each approach with varying number of patient samples in the training data. IC50 summary statistics were not used because only one cell line was a non-responder according to this summary statistic.

When using at least 60 patients for training, the best P2P method performed significantly better than C2P methods for AUC and Slope binarized response statistics (p = 0.003 and 1.2e-07 for AUC and Slope respectively, using one-sided Mann-Whitney tests). CP2P-AUC using 60 patients and CP2P-Slope using 80 patients performed significantly better than P2P using 100 patients (p = 0.004 and 0.007 respectively using one-sided Mann-Whitney tests, Figure 5C). Consistent with the other drugs, these results indicate that once samples from enough patients become available, the patient-trained classifiers outperform C2P classifiers, i.e., the predictive value of classifiers using cell lines alone is limited. Yet, classifiers using both cell lines and patients outperform solely patient based classifiers with only a fraction of the patient samples needed for patient-based methods.

## Discussion

We have shown that models developed with both cell lines and patient samples can predict drug response as well as or better than those using cell line or patient samples alone. The best performing CP2P models performed significantly better than the best performing C2P and P2P models for three out of four tested drugs, and in case of the fourth tested drug, CP2P and P2P did not statistically significantly differ in performance. The real difference in performance became apparent when more than 30 patients were available at training time. Even when more than 100 patient samples are available, the best CP2P models still significantly outperformed the best P2P models.

For docetaxel and erlotinib, we were able to compare performance of the models using cell lines across many tissue types to those matching the tissue type of the patient’s cancer. For docetaxel, both C2P and CP2P performed better when all cell line samples were used. For erlotinib, classifiers using all cell lines performed similarly to lung cancer-only classifiers, though the variance of the lung cancer-only classifiers was lower. We thus conclude that using all cell lines is beneficial in cases when there is only a small number of matching-tissue cell lines. Further experiments are needed to make definitive conclusions in cases when large numbers of tissue-matching cell lines are available.

Performance of the models combining heterogeneous sources of data often hinges on the pre-processing step we call homogenization. In the case of bortezomib, for example, patients’ mRNA expression was observed to be orthogonal to the expression of cell lines from the CGP dataset when projected onto the first two principal components. We have chosen to address this problem by using ComBat and SVA. However, this approach prevents easy application of our prediction models for single samples originating from heterogeneous sources. Single-sample batch effect correction methods are currently under active development ^29^; they have the potential to boost the performance of both C2P and CP2P models in future applications.

Analyzing the performance of the range of classifiers we used in our experiments, we note that no single model outperformed others across all drugs. In fact, methods that performed significantly better for some drugs (*SVM lin all* in bortezomib with binarized Slope as the response summary statistic), had significantly worse performance for others (epirubicin). Interestingly, models trained with whole genomes and mRMR1000 tend to do better than the ones trained with L1000 genes across the tested clinical trials. Gene selection (mRMR) seems to be of more importance when the number of available patients is small (erlotinib and docetaxel). Similarly to other studies ^18,24,25^, we show that for different drugs, different measures of response (IC_50_, AUC, Slope) appear to yield the best predictors. These results highlight the importance of the selection of a consensus measure for development of genomic biomarkers of therapy response.

In conclusion, our extensive experiments highlight the importance of data integration across *ex vivo* and *in vitro* data to achieve the best performance in drug response prediction. Furthermore, when the number of patients available for training is sufficient, CP2P models are able to achieve as good of a performance as models trained using solely patient data while requiring substantially fewer patients. Our results therefore support the relevance of preclinical data for therapy prediction in clinical trials, motivating more efficient and cost-effective trial designs.

## Methods

The overall design of our study is represented in Figure 1.

### Data Preprocessing

The CGP cell line dataset was pre-processed as in Haibe-Kains et al. ^18^. We used fRMA ^30^ to preprocess the RAW CEL files, and then applied Jetset ^31^ to select the optimal probe set for each gene. The cell lines from Heiser ^28^ were processed the same way. Pre-processed BATTLE lung cell line samples were available from GEO. The RAW CEL files for erlotinib’s clinical dataset was downloaded and preprocessed using RMA ^32^, as implemented by the *rma()* function in the *affy* package ^33^, and custom CDF files ^34^ (version 19) was used to map probe sets to the ENTREZ gene IDs. If more than one probe set was matched to a gene symbol, the mean of the expression values was used. For other clinical datasets, the preprocessed data was used as downloaded from GEO (Supplementary Datasets section). The PCA plots were created using vqv package ^35^. The ellipses represent one standard deviation away from the mean of the Gaussian fitted to each data type.

### Homogenizing cell line and patient datasets

For C2P and CP2P, the intersection of the genes among cell line and patient datasets was used. We first attempted to homogenize the data with ComBat. If PCA analysis showed that a considerable amount of orthogonality between the patient and cell line datasets remained, SVA was used instead. For bortezomib, SVA with number of surrogate variable set to 3 was first applied to the cell line mRNA expression data. We then applied SVA to cell line and patient data combined with the number of surrogate variable set to 2; for docetaxel and erlotinib, ComBat was used; for epirubicin, SVA was used with the number of surrogate variables set to 3.

### Classifier description

In our study, we used seven classifiers based on diverse machine learning approaches. Ridge logistic regression (labeled as *Ridge*) is a regression model, assigning weight to each feature to make a binary prediction. L2 regularization shrinks the weights to avoid overfitting. Lasso logistic regression (labeled as *Lasso*) ^36^ works the same way as Ridge but uses L1 regularization instead, which sets some of the weights to zero, effectively performing feature selection again to avoid overfitting. Elastic net logistic regression (labeled as *Elastic Net*) ^37^ also works similarly to Ridge but uses a linear combination of L1 and L2 regularizations and is able to select correlated features (through L2) while still performing feature selection (setting some of the weights to zero) through L1. Random forest (*RF*) ^38^ uses an ensemble of decision trees to make classification predictions. Each decision tree uses a random subset of features trained on a bootstrapped set of samples. The output is the mode of the classification from all decision trees in the random forest. Support Vector Machines (*SVM*) ^39^ make classification predictions by first transforming the data according to a chosen kernel and then constructing a maximum margin classifier such that the different classes are separated by the decision hyperplane as much as possible. SVM with linear kernel (labeled as *SVM lin*) performs a linear transformation of the data, whereas SVM with radial basis function kernel (labeled as *SVM rbf*) performs a Gaussian transformation of the data. Similarity Network Fusion with label propagation (labeled as *SNF*) ^40^, constructs a similarity network on all the samples and uses label propagation ^41^ to make classification predictions given the labels of the training set.

Prior to training any of the classifiers, we used three settings for constraining feature space (genome-wide gene expression profiles) by means of feature selection: L1000, mRMR1000, and all genes. The L1000 genes ^42^ a set of 1000 genes that have been carefully chosen and are able to capture approximately 80% of the information in the human genome. mRMR ^23,43^ is a feature selection algorithm that constructs a set of features with minimal redundancy to each other and maximal relevance to the given label. mRMR selection was made using the training set.

### Summary of the drug dose-response curves in cancer cell lines

It has been previously shown that different summary statistics of the drug dose-response curves have varying degree of reproducibility ^18^ and relevance in predicting patients’ therapy response ^24,25^. To analyze this effect, we compared three binary response/non-response labels derived from three different summary statistics of drug response for cell lines: the concentration required to inhibit 50% of the maximal cell growth (IC_50_), the area under and the slope of the drug dose-response curve (referred to as AUC and Slope, respectively, for details please see Supplementary Methods). We ran all experiments for C2P and CP2P separately for each binarized outcome denoting results with the suffixes “-IC50”, “-AUC”, and “-Slope”.

### Training and testing of the classifiers

For P2P and CP2P experiments, we varied the number of patient samples in the training set. Any patient sample not in the training set was used in the test set. For example, for bortezomib’s P2P experiments, we varied the number of patient samples used for training from 20 to 150 at an increment of 10, while for C2P experiments, the test set consisted of all patients. The training set for P2P consisted of patient samples only, for C2P -- cell line samples only, and for CP2P -- all available cell line samples with the corresponding portion of the patient samples.

For P2P, it was also necessary to ensure that at least 5 samples from each class (responder/non-responder) were present in both training and test sets. This ensured that training was possible, as at least a few examples from both labels are necessary to build a model. After partitioning the data into train and test sets, 5-fold CV was used on the training set for the model parameter selection (e.g. the strength of the L1 regularization in Ridge, or the number of decision trees used in Random Forest) and for training of the models. AUROC requires at least 10 samples to be meaningful, therefore if the CV set had fewer than 10 samples, the model parameters were optimized for accuracy; otherwise, parameters were optimized for AUROC.

We used 7 classifiers and 3 different feature selection settings, yielding 21 different models. We used 3 different binarized cell line response summary statistics for C2P and CP2P each resulting in 6 different training scenarios. In the P2P scenario we used the response/non-response labels obtained as discussed above for each specific clinical trial, resulting in one training scenario for P2P. The total number of model-outcome-training type models is then 21 * (6 + 1 (for P2P)) = 147. For erlotinib, IC_50_ and AUC summary statistics produced the same drug response labels, so the effective number of tested models was 105. For epirubicin, IC_50_ summary statistics produced highly unbalanced labels and therefore was not used resulting in a total of 105 model-scenario comparisons.

### Research reproducibility

All analyses were performed in R version 3.1.1. We used the glmnet ^44^ package for Elastic Net, Lasso, and Ridge; the randomForest ^45^ package for RF, the kernLab ^46^ package for SVM, the SNFtool package ^47^ for SNF, and the mRMRe package ^43^ for mRMR. Training of RF and SVM was done using the caret package ^48^. The AUROC values were calculated using the ROCR ^49^ package. Our experiments are fully reproducible (see more in the Supplementary Information). The code and the RData are available at http://compbio.cs.toronto.edu/cp2p/.

## Footnotes

1 also referred to as companion tests

2 Note that training of the breast tissue only cancer cell line models for docetaxel was done using 3-fold cross validation due to the small number of tissue-specific cell lines.

## Abbreviations

AUC: Area under the drug dose-response curve
AUROC: Area Under the Receiver Operating Characteristic Curve
C2P: Model predicting patients’ drug response from in vitro (cancer cell lines) data
CDF: chip definition file
CGP: Cancer Genome Project
CP2P: Model predicting patients’ drug response from the combination of in vitro (cancer cell lines) and ex vivo (patient tumors) data
IC50: Drug concentration required to inhibit 50% of the maximal cellular growth of a given cell line
NSCLC: non-small cell lung cancer
P2P: Model predicting patients’ drug response from ex vivo (patient tumors) data
PCA: principal component analysis
ROC: receiver operating characteristic

## Authors’ contributions

CZ wrote the code, performed the analysis, and drafted the paper. YL wrote the code and prepared the data. AG and BHK conceived of the study, supervised it and wrote the paper. BHK and ZS collected and processed the pharmacogenomic data. All authors edited and approved the final manuscript.

## Acknowledgments

The authors would like to thank CCSRI Innovation Grant 703471. The authors thank the Natural Sciences and Engineering Research Council of Canada, University of Toronto, Sick Kids Foundation, Cancer Research Society and the Gattuso Slaight Personalized Cancer Medicine Fund at Princess Margaret Cancer Centre for their generous funding.

